# namiRa: A Comprehensive, Manually Curated Database for MicroRNA Expression, Function, and Deregulation in Cancer

**DOI:** 10.1101/2025.11.25.671411

**Authors:** Saeed Mohebbi, Alireza Dostmohammadi, Sara Amjadian, Zahra Abdi, Hanieh Torkian, Mojtaba Ghavidel, Mohadese Rahbar, Afsaneh Yazdani Movahed, Niloofar Rastidoust, Forouzan Mahdizad, Azam Akbari, Shamimeh Mosanan Farsi, Fatemeh Azadedel, Leila Abdollahi, Raha Farhadnejad, Nafiseh Salmanzadeh, Hanieh Sadeghi, Sharif Moradi

## Abstract

MicroRNAs (miRNAs) have the potential to serve as oncogenes or tumor suppressors, playing important roles in the pathogenesis of human cancers. Despite the growing recognition of miRNA significance, the contributed information remains scattered in the publications. This fragmentation underscores the need for a centralized and comprehensive database that consolidates miRNA expression patterns, functional roles, and regulatory interactions across diverse cancer types. So far, several miRNA databases have been developed, but neither of them enables in-depth functional analysis or comparative visualization of miRNA data. Here, we present namiRa, a manually curated database, to offer a comprehensive resource for miRNA expression and functional significance in various types of cancer, describing miRNA-cancer associations based on a thorough review of the literature. namiRa provides an extensive collection of miRNA expression profiles, detection methods, functional analyses for miRNAs *in vitro* and *in vivo*, and visualized regulatory networks across different cancer types. The current version of namiRa documents curated relationships between 1,077 human miRNAs and 33 types of human cancers, based on data from 9,884 published papers. namiRa is accessible at https://www.namira-db.com.

## Introduction

MicroRNAs (miRNAs) are short (∼22-nucleotide long) regulatory noncoding RNAs that silence gene expression at the posttranscriptional level via binding to the 3′ untranslated region (UTR) of the targeted mRNAs [1, 2]. miRNAs are expressed in virtually all cells and play crucial roles in regulating various biological processes such as cell survival, proliferation, differentiation, migration, and apoptosis [3–6]. Therefore, alterations in miRNA expression and function are associated with aberrant physiological conditions and many human diseases, particularly cancers [7–10]. Moreover, genome-wide association studies indicate that a large number of human miRNA genes are frequently located at genomic regions involved in cancer [11, 12]. The dysregulation of miRNAs has been shown to promote or repress the cancer phenotype. Nevertheless, the field still lacks a comprehensive repository that consolidates miRNA expression data with their corresponding functional annotations and regulatory network interactions across the spectrum of human cancers.

In recent years, with the increasing number of identified miRNAs, detailed information on miRNA–cancer associations remain dispersed across the literature [13]. Researchers often face challenges in integrating data from multiple sources to gain a holistic view of miRNA dysregulation in cancer. Hence, dozens of online miRNA-related resources have been developed to facilitate the study on miRNA functions and regulatory mechanisms. miRBase is the primary miRNA sequence repository that provides the sequence, annotation, and nomenclature of known miRNAs from different species [14]. miRTarBase [15] and miRecords [16] are notable for their extensive collections of experimentally validated miRNA-target interactions. miRWalk [17], miRDB [18], and TargetScan [19] present computationally predicted miRNA-target relationships, offering insights into potential regulatory networks. MSDD collects and integrates experimentally supported disease-associated miRNA single-nucleotide polymorphisms [20]. miRCancer employs a text-mining approach to compile miRNA expression states in cancer cells [13]; however, it provides no information regarding the functional and mechanistic significance of various miRNAs in cancer. Likewise, HMDD [21], RNADisease [22], OncomiR [23], and OMCD [24] databases explore the miRNA expression levels qualitatively or quantitatively without direct functional evidence. miR2Disease offers a resource for associations of miRNAs with different diseases, extracted from peer-reviewed research articles [25]; however, the data that it provides for various miRNA-cancer relationships is very limited or superficial. dbDEMC [26] and CMC [27] present the functional annotations of miRNAs associated with cancer hallmarks based on bioinformatic analysis. While experimentally verified miRNAs with functional patterns are annotated in OncomiRDB [28], the number of included miRNAs (300) is relatively limited. Therefore, a comprehensive resource that integrates miRNA expression patterns, their experimentally validated functional roles, and regulatory interactions across diverse cancer types has remained a significant gap in the field. To bridge this gap, we have developed a manually curated database entitled namiRa, which provides an extensive resource for miRNA regulation and deregulation in cancer. namiRa (which means ‘immortal’ in Persian, reflecting on the immortality of cancer cells) is the most comprehensive database dedicated to cataloging miRNAs in various human cancers. The most important and unique features of namiRa include: (i) a comprehensive resource for representing the miRNA expression patterns and detection methods in different human tumors and cancers; (ii) a repository for the functional annotations and experimentally validated outcomes of miRNA dysregulation in cancer; (iii) bridging miRNA expression data and the target gene databases through hyperlinking; (iv) presentation of the detailed results of more than 9,884 published papers on miRNA-cancer relationships; and importantly (v) dynamic illustration of a unique miRNA-gene interaction schematic map of the regulatory networks for each individual miRNA species in each cancer type. This visualization of the miRNA regulatory processes offers a valuable and quickly accessible information on the most important, experimentally verified interactions of the miRNA, its target genes and affected cellular processes. namiRa will assist researchers in comparative studies between different miRNAs in different cancers, experimental designs and verification, understanding involved mechanisms in cancer, and clinical investigations.

## Methods

### 1. Data collection and database content

namiRa aims to provide a web-based data resource for the search and analysis of experimentally verified miRNA-cancer associations. To create namiRa, a comprehensive search of the PubMed database was conducted for publications from 2000 to 2023, employing keywords such as “microRNA cancer”, “miRNA tumor”, “miRNA AND cancer”, and “microRNA tumour” within the full text. More specifically, the term “microRNA” or “miRNA” was searched alongside the name of a desired tumor/cancer type, such as “microRNA” AND “gastric cancer”. From the reported results, non-relevant and review articles were excluded. Moreover, since namiRa focuses on the impact of miRNAs on cellular behaviors in cancer, papers lacking experimentally supported functional analyses of miRNAs were excluded. We also excluded studies based solely on miRNA expression analysis in bodily fluids (*e.g.,* serum, plasma), as the database emphasizes functional data in a tissue/cellular context. An exception was made for all types of blood cancers, which were included irrespective of the sample source.

Thereafter, a table was designed in which each entry contained the detailed information on the miRNA-cancer associations, including cancer type, miRNA ID, expression pattern of the miRNA correlated with each cancer type, the detection method used to determine the miRNA expression pattern, the upstream regulators of miRNAs investigated in the articles, and a description of the miRNA function in promoting or inhibiting of each cancer type based on *in vitro* and *in vivo* studies. In addition, experimentally validated target genes were extracted from the corresponding literature references. In this regard, we included only those gene targets that had been fully verified as direct and functional gene targets, meaning that we ignored those targets which were just downregulated upon miRNA overexpression without additional experimental support. To ensure consistent data interpretation, all experimentally observed miRNA effects were standardized and reported as the functional outcome of increased miRNA expression. For instance, if the suppression of a miRNA was found to inhibit cancer cell proliferation, this was interpreted and recorded in the database as the miRNA having a pro-proliferative role when overexpressed. Conversely, if miRNA suppression promoted apoptosis, the miRNA was annotated as anti-apoptotic. As the other rule, we integrated the miRNA information from several articles that were in the same direction of associations, either increased expression and oncogenic properties or decreased expression and tumor suppressor properties. Regarding the miRNA arms, we selected the strand with the higher read number in miRBase database as a functional arm (*i.e.* the guide strand) if no distinction was made in the papers, and nothing was specified if there was no record in the database. To construct the content of namiRa, we meticulously reviewed and included the detailed information from more than 9,884 relevant publications. Each article was carefully evaluated to extract data on miRNA-cancer associations, following a rigorous process to ensure the inclusion of high-quality and reliable information. From this extensive review, we documented associations between 1,078 human miRNAs and 33 distinct cancer types.

In the namiRa database, we also designed interactive diagrams of miRNA regulatory networks to visually illustrate the main validated target genes and the cancer-driving cellular processes affected by miRNAs. Furthermore, convenient links to other miRNA databases have been offered by namiRa, including miRBase, TargetScan, miRDB, TarBase [29], and miRTarBase which present experimentally validated miRNA target genes. A link to the mirPath database [30] is also provided for each miRNA, enabling users to explore the involvement of the miRNA in cellular pathways and processes. Figure 1 shows the database preparation procedure.

**Fig. 1.**
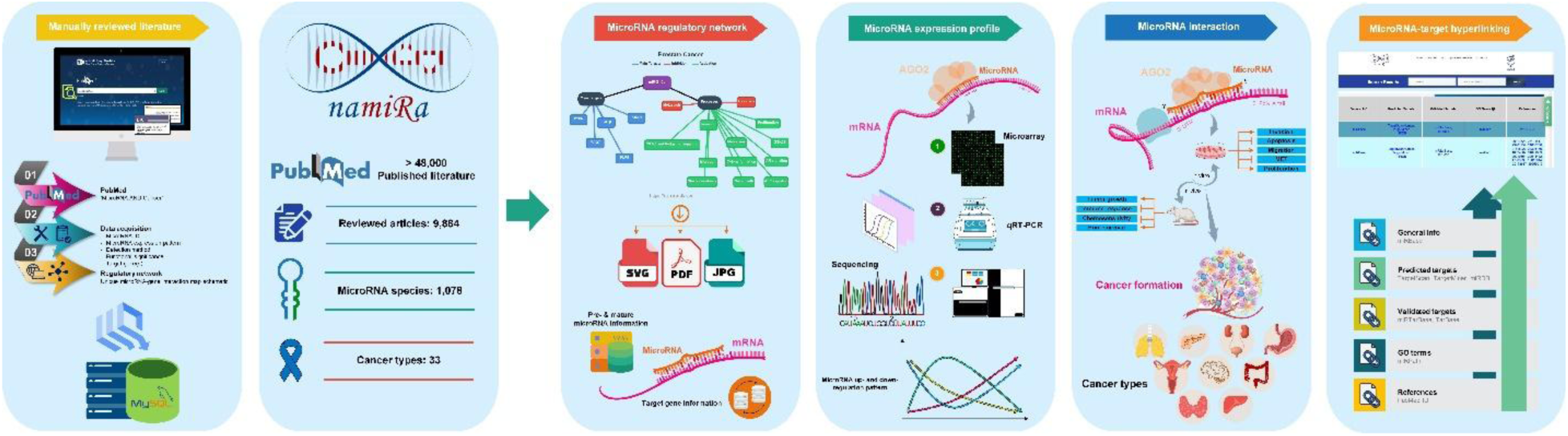
Schematic overview of the namiRa database architecture. The schematic illustrates the core features of namiRa, including: (1) comprehensive coverage of miRNA species and associated literature, (2) cancer type-specific miRNA regulatory networks, (3) miRNA expression profiles, (4) functional annotation of cancer-related miRNAs, (5) links to miRNA-target interactions in external databases, and (6) general miRNA information, including the miRNA-associated cellular processes.

### 2. Database organization and web interface

The namiRa database is maintained on powerful servers with high bandwidth and configurations to support simultaneous usage by multiple researchers. We provide an SSL connection to ensure secure access to the namiRa database. The database runs on Apache 2.4.57 as the web server. The backend of the database is written in PHP 7.7, and the data is stored on a MySQL 5.7.43 server. The frontend has been developed using HTML5, CSS3, JavaScript, and jQuery. Additionally, Ajax is implemented for live searches. To enhance functionality and usability for researchers, we have developed an API in both R and Python for direct access to the database.

## Results

### 1. Database usage: Search page

namiRa provides a search engine to query detailed miRNA-cancer associations documented in the database. Users can search using a miRNA ID, a cancer type, or both, allowing for independent or combined (AND operation) queries (Figure 2). The search engine features live search functionality, offering dynamic suggestions as users start typing. For instance, entering a few letters of a miRNA ID or cancer type generates a list of matching entries, making it easier for users to locate relevant terms available in namiRa. To enhance flexibility, namiRa supports searches using terms that align with documented cancer types. For example, if a user searches for “Hepatocellular Carcinoma” instead of “Liver Cancer,” the system can still retrieve results for “Liver Cancer” based on a predefined list of frequently used synonyms. While this feature currently supports a limited set of terms, namiRa plans to expand its coverage in the future for even more flexible search options.

**Fig. 2.**
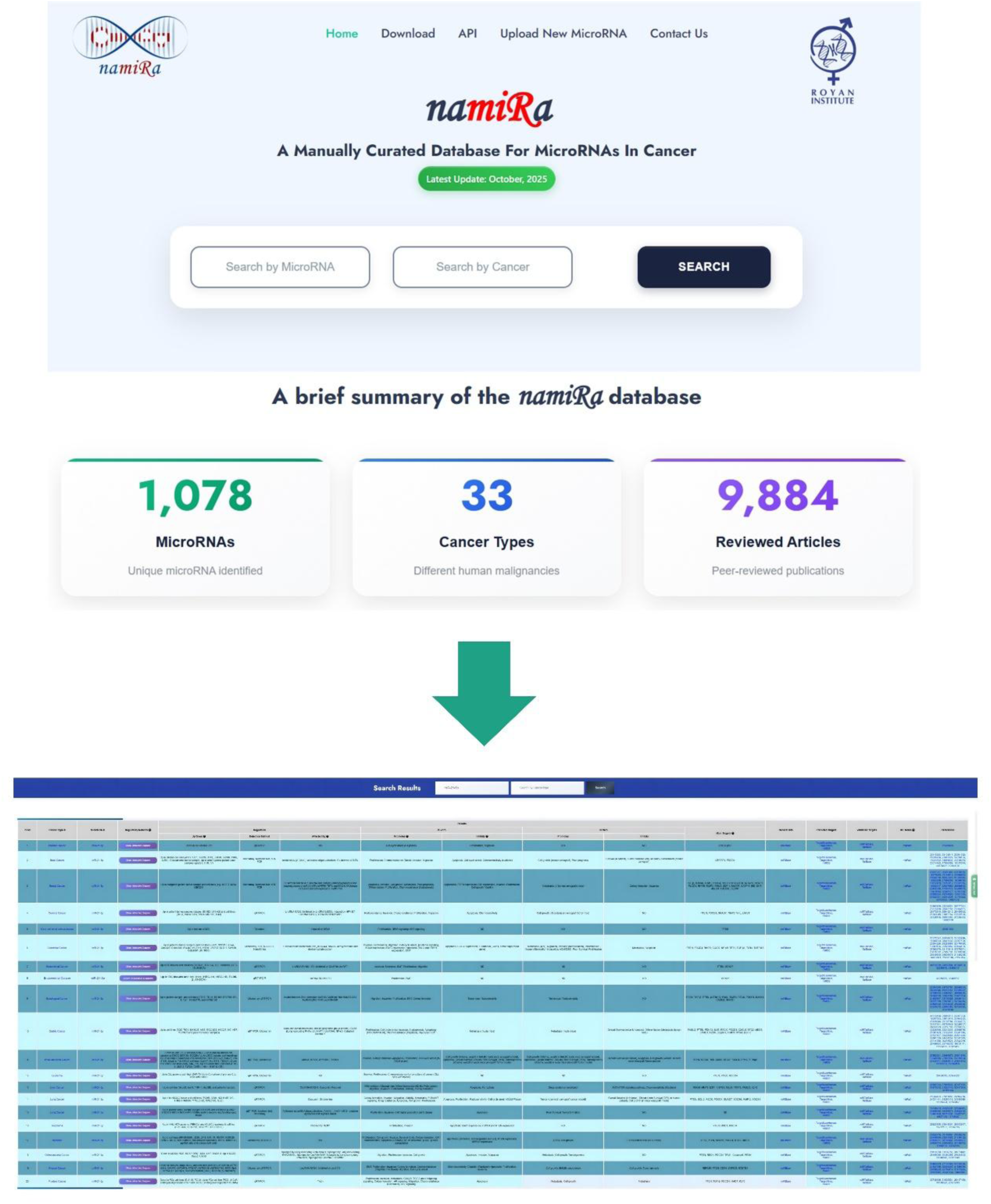
The query workflow of the namiRa database. The schematic depicts the search interface, including dynamic query options (miRNA/cancer terms), autocomplete suggestions, literature citations, and links to external miRNA resources (target predictions, validated miRNA targets, and ontologies). Designed to accelerate research into miRNA-cancer interactions, the workflow streamlines the exploration of miRNA-cancer associations by integrating curated data with external resources.

To improve the robustness and user-friendliness of the search functionality, namiRa incorporates a semantic search feature for cancer names. This system leverages the OpenAI large language model GPT-OSS-120B [31], accessed via GroqCloud API services (https://groq.com/groqcloud), to capture the conceptual meaning of user queries. Consequently, if a user enters a cancer name that is conceptually similar but not an exact textual match to the curated terms in the database, the system can intelligently retrieve the relevant results. This capability ensures that researchers can successfully find pertinent data even when using synonyms or related terminology, thereby reducing missed queries and enhancing the efficiency of data retrieval.

Similar to searching by cancer terminology, miRNA nomenclature can be complicated and problematic. Because original publications may lack sufficient information on the exact miRNA species within a large family or may not distinguish between different mature miRNA arms, namiRa’s flexible search function allows for miRNA queries without requiring exact identification data. Thus, if a query returns multiple miRNA IDs, users can select the specific miRNAs of interest. After entering a query - whether a miRNA name, cancer type, or both - or selecting from the live search suggestions, users can click the search button to view the results.

### 2. User interfaces: Results page

The results table (Figure 2) allows for sorting entries by “Cancer Type” or “MicroRNA” in ascending or descending order. Additionally, users can refine their search directly on the results page without returning to the search page. Clicking on a specific cancer type or miRNA ID within a row in the results table takes users to a detailed page, displaying all information related to that particular entry (Figure 2). This seamless navigation allows users to efficiently explore the miRNA-cancer relationships documented in the database. On the opened result page, miRNA information has been provided in both graphical and text-based interfaces as discussed below.

#### 2.1. Graphical interface

namiRa provides a dynamic graphical interface for each miRNA and cancer type to illustrate validated miRNA targets and their associated cancer-related cellular processes in the form of an interactive network. This interface, titled “Interactive Regulatory Network,” is generated on a new page by clicking the “show interactive diagram” button and can be downloaded in SVG, JPG, and PDF formats. Within the network, activated or inhibited cellular processes are color-coded to clarify the functional mechanisms of the miRNAs. Moreover, each node in the network can be repositioned, allowing users to categorize data visually based on custom criteria.

#### 2.2. Text-based interface

In its text-based interfaces, namiRa presents annotated miRNA information under three main categories: miRNA expression patterns, functional mechanisms, and hyperlinks to external resources, each of which is described below.

##### 2.2.1. The miRNA expression panel

Increasing evidence has shown that dysregulation of miRNA expression contributes directly to cancer initiation and progression, as miRNAs can function as tumor suppressors or oncogenes [32]. Tumor-suppressive miRNAs are typically downregulated in tumorigenesis, whereas oncogenic miRNAs are overexpressed [33]. Accordingly, we examined the expression patterns of numerous miRNAs across different tumor tissues and cell types from the literature. We cataloged more than 5,000 miRNA entries across 33 cancer types. Among these, 65.6% (range: 50.0%-77.1%) were downregulated tumor-suppressive miRNAs, and 26.9% (range: 16.6%-41.6%) were upregulated oncogenic miRNAs. Notably, an average of 7.3% of the reported miRNAs showed context-dependent expression, being either over- or under-expressed depending on the condition. Given the spatiotemporal specificity of miRNA expression, it is not surprising that some miRNAs exhibit different patterns in different contexts or even in different cell types within the same context [34]. Therefore, further experimental evidence is required to clarify the expression direction of such miRNAs. Table 1 summarizes the percentages of upregulated, downregulated, and context-dependent miRNAs for each cancer type.

**Table 1.** The percentage of up- and downregulated miRNAs in each cancer type.

| **Cancers\*** | **Upregulated miRNAs (%)** | **Downregulated miRNAs (%)** | **Up/Downregulated miRNAs (%)** |
| --- | --- | --- | --- |
| Brain cancer | 27.19 | 69.29 | 3.50 |
| Head and neck cancer | 16.66 | 69.50 | 13.82 |
| Eye cancer | 27.18 | 70.87 | 1.94 |
| Thyroid cancer | 28.12 | 67.50 | 4.37 |
| Lung cancer | 21.49 | 66.60 | 11.90 |
| Breast cancer | 37.05 | 57.58 | 5.35 |
| Kidney cancer | 25.84 | 67.79 | 6.36 |
| Bladder cancer | 24.44 | 68.88 | 6.66 |
| Esophageal cancer | 34.43 | 62.26 | 3.30 |
| Stomach cancer | 19.74 | 77.17 | 3.07 |
| Colon cancer | 28.30 | 61.72 | 9.97 |
| Liver cancer | 38.75 | 61.25 | 0.00 |
| Pancreatic cancer | 27.73 | 64.06 | 8.20 |
| Uterine cancer | 28.08 | 65.73 | 6.17 |
| Cervical cancer | 20.07 | 71.96 | 7.95 |
| Ovarian cancer | 21.64 | 68.04 | 10.30 |
| Prostate cancer | 21.08 | 70.06 | 8.84 |
| Blood cancer | 26.60 | 55.96 | 17.43 |
| Bone cancer | 21.10 | 70.10 | 8.29 |
| Soft-tissue cancer | 48.00 | 52.00 | 0.00 |
| Skin cancer | 28.29 | 62.43 | 9.26 |

\*Specific cancer types: Retinoblastoma (Eye cancer); Clear cell renal cell carcinoma and renal cell carcinoma (Kidney cancer); Endometrial cancer and uterine leiomyoma (Uterine cancer); Leukemia, lymphoma, and myeloma (Blood cancer); Osteosarcoma and Ewing sarcoma (Bone cancer); Ewing sarcoma and fibro-sarcoma (Soft-tissue cancer); Cutaneous basal cell carcinoma, cutaneous hemangioma, cutaneous squamous cell carcinoma, cutaneous T cell lymphoma, eyelid sebaceous gland carcinoma, melanoma, and Merkel cell carcinoma (Skin cancer); Brain cancers (Brain cancer); Prostate cancer (Prostate cancer); Hepatocellular carcinoma (Liver cancer); Ovarian cancer (Ovarian cancer); Cervical cancer (Cervical cancer); Gastric carcinoma (Stomach cancer); Head and neck cancer (Head and neck cancer); Pancreatic cancer (Pancreatic cancer); Colon/colorectal cancer (Colon cancer); Bladder cancer (Bladder cancer); Esophageal adenocarcinoma/squamous cell carcinoma (Esophageal cancer); Lung cancers (Lung cancer); Thyroid cancer (Thyroid cancer); Breast cancers (Breast cancer).

Following the determination of expression patterns, the methods used for miRNA detection and the upstream regulators of miRNA expression are also detailed in separate columns. The detection strategies for deriving miRNA expression patterns included qualitative methods, such as northern blot and in situ hybridization (ISH), and quantitative methods, such as high-throughput sequencing and, most frequently, qRT-PCR. Additionally, a column entitled “Affected by” reports the upstream miRNA regulators identified in the literature, including long non-coding RNAs, circular RNAs, epigenetic modifiers, signal transduction inhibitors, and various drugs, particularly chemotherapeutic agents.

##### 2.2.2. The miRNA functional mechanism panel

miRNAs are involved in all aspects of cancer biology, including cell proliferation, migration, invasion, apoptosis, and angiogenesis. Based on their target genes, miRNAs can function as oncogenic or tumor-suppressive RNAs [35]. To facilitate the study of corresponding miRNA functions and regulatory networks in cancer, we collected detailed, direct functional evidence for each miRNA’s role in either promoting or inhibiting cancer, based on *in vitro* and *in vivo* studies. We cataloged miRNAs that regulate at least one cancer-attributed cellular process in cell cultures or animal models. Consistent with the miRNA expression data, an average of 23.81% of miRNAs were positively associated with cancer hallmarks, whereas 63.88% suppressed cancer development, suggesting a predominantly tumor-suppressive role for the miRNA biogenesis pathway. The functional role of miRNAs can also be context-dependent due to the regulation of various target genes and signaling pathways [34]. Accordingly, we found that 12.25% of miRNAs have dual and distinct functions in specific cancer types or under specific conditions (Figure 3a). Moreover, the greatest number of miRNAs investigated *in vitro* were detected in brain and thyroid cancers, followed by eye, kidney, bladder, and blood cancers (with soft-tissue sarcoma excluded due to limited publications). In contrast, the greatest number of miRNAs studied in both *in vitro* and *in vivo* models were associated with colon, pancreatic, stomach, liver, and bone cancers. Figure 3b shows the distribution of miRNAs reported in *in vitro* studies only versus those validated in both *in vitro* and *in vivo* models.

**Fig. 3.**
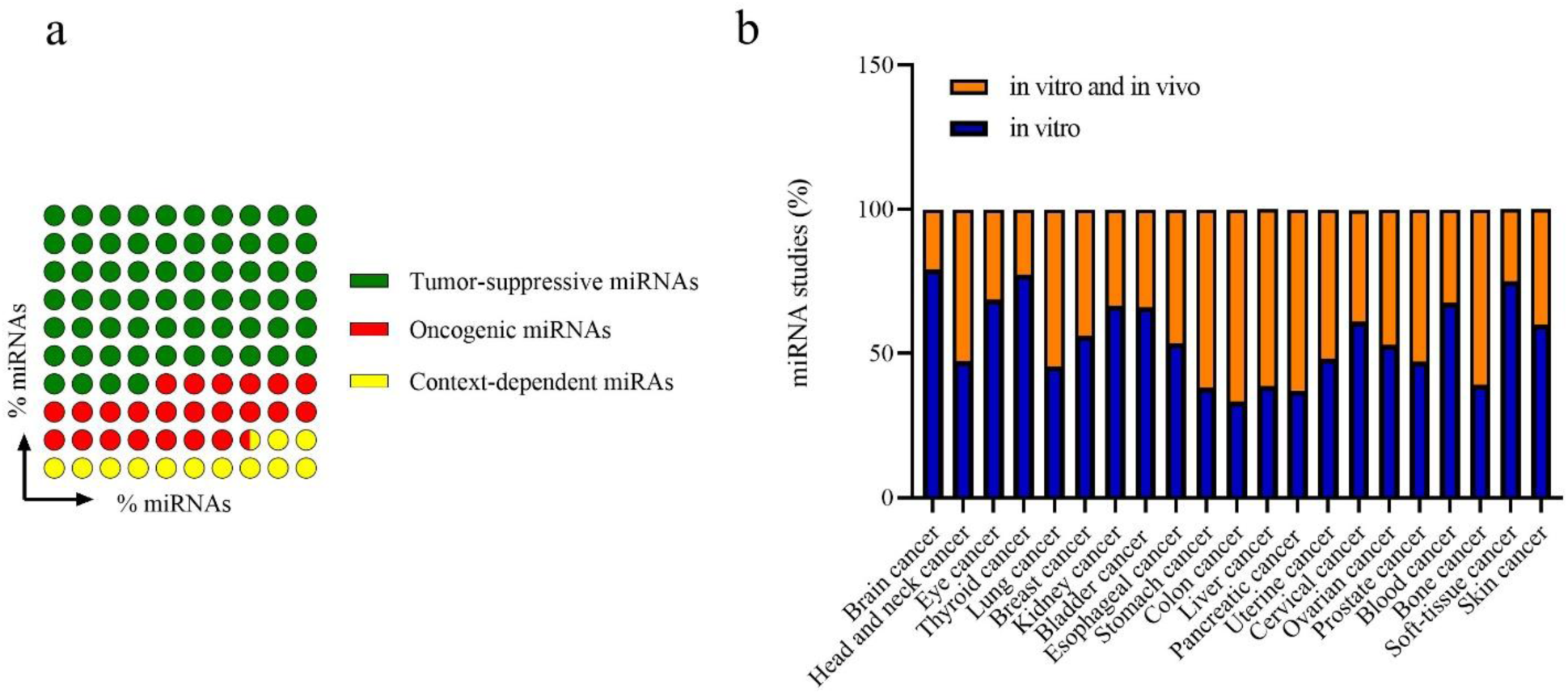
Functional characterization of miRNAs in cancer based on *in vitro* and *in vivo* studies. (a) Pie chart showing the proportion of miRNAs with tumor-suppressive (63.88%), oncogenic (23.81%), or context-dependent (12.25%) functions. (b) Bar plot showing the distribution of miRNAs validated in *in vitro* studies only versus those validated in both *in vitro* and *in vivo* models across different cancer types. Brain and thyroid cancers had the highest number of miRNAs studied *in vitro*, whereas colon, pancreatic, stomach, liver, and bone cancers had the highest number of miRNAs validated in both *in vitro* and *in vivo* models.

In another column of the functional mechanism panel, we also provide the main experimentally verified target genes for each miRNA, as validated by robust methods in specific cancer types. The largest numbers of target genes were annotated for miR-145-5p, miR-34a-5p, miR-124-3p, and miR-20a-5p.

##### 2.2.3. Hyperlinks to external databases

To facilitate a deeper understanding of miRNAs, their target genes, and involved pathways, namiRa provides direct access to other miRNA databases via hyperlinks that open in new pages. Basic miRNA information can be acquired from miRBase, an archive of miRNA sequences and annotations. Links to TargetScan Human, TargetMiner, and miRDB provide access to predicted miRNA target genes, while hyperlinks to miRTarBase and TarBase lead to databases of experimentally validated miRNA-gene interactions. miRPath is also included as a web server for miRNA pathway (de)regulation analysis. Furthermore, reference publications are accessible via PubMed IDs on the database’s main page.

It is worth noting that in the search results, categories containing the query terminology are highlighted in bold, and a hyperlink is generated for each category with documented miRNA-cancer relationships. These hyperlinks allow users to retrieve all miRNA entries associated with the selected cancer terminology.

##### 2.2.4. Integration of an AI assistant for enhanced data exploration

To further facilitate user interaction and data interpretation, we integrated an AI-powered assistant directly into the namiRa platform. This feature, accessible from the search results page, is built upon the OpenAI GPT-OSS-120B large language model [31] to provide users with an intuitive, conversational interface for querying the database’s knowledge domain. Researchers can utilize this tool to ask specific questions about miRNAs or cancer types, investigate potential mechanistic links between them, or pose general inquiries related to miRNA biology in cancer. The assistant is designed to synthesize information and offer immediate, context-aware insights, complementing the structured data presentation. To ensure sustainability and consistent performance for all users, the service operates freely within defined usage limits. The underlying model, its prompting instructions, and the interface text remain adaptable, allowing for future refinements to maximize the accuracy and utility of the generated responses for the research community.

### 3. Submit page

namiRa provides an ‘**Upload New MicroRNA**’ page that allows researchers to submit information from published articles on miRNAs related to cancer that have not been previously documented in the database. The submitted records will be included in the database and made available for users and researchers after approval by the submission review committee.

## Discussion

In this study, we developed namiRa, an extensive database that provides detailed information on the oncogenic or tumor-suppressive roles of miRNAs in various cancers. It serves as a centralized tool for studying the expression, functions, and interactions of a large number of miRNAs, enabling cancer researchers to retrieve and analyze data more comprehensively. namiRa is a repository of experimentally validated miRNA data without additional computational processing, allowing users to perform analyses based on their own criteria.

Within the archive, we identified the cancers with the most annotated miRNA entries as lung (600), bone (401), stomach (391), and colon (380), respectively (Figure 4). Moreover, miR-21-5p, which is a well-known oncogenic miRNA involved in a majority of cancer types, was the most frequently studied. miR-34a-5p and miR-145-5p were also among the most experimentally investigated miRNAs, followed by miR-200s-3p and miR-205-5p. Table 2 shows the most studied miRNAs for each specific cancer.

**Fig. 4.**
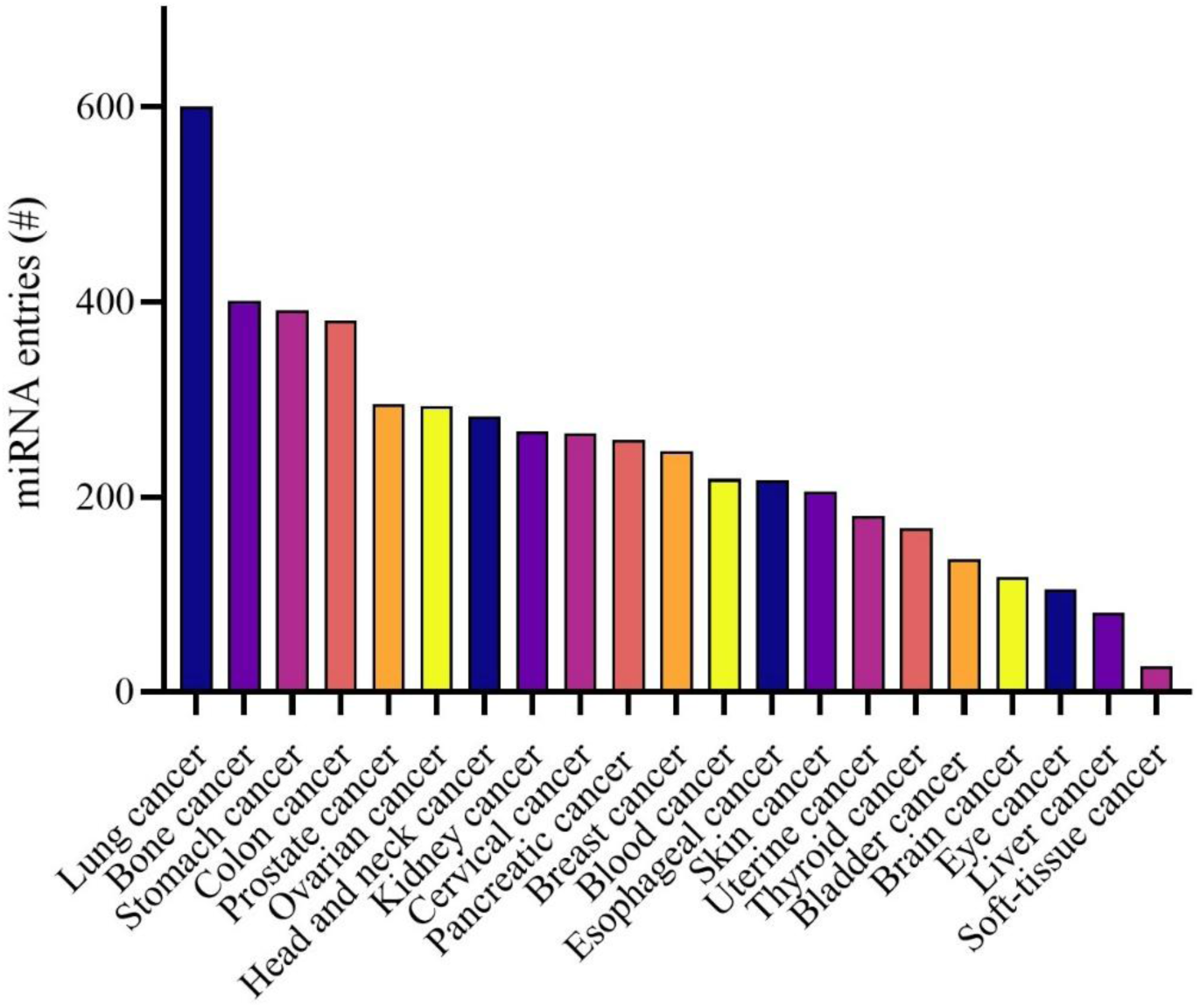
Cancers annotated for miRNA investigation. Twenty-one major cancer types, encompassing 33 specific cancer (sub)types, were examined for miRNA data collection through an extensive literature review. Lung, bone, stomach, and colon cancers had the highest number of miRNA entries, respectively.

**Table 2.** The most studied miRNAs in each cancer.

| Cancer type | The most experimentally investigated miRNAs |
| --- | --- |
| Brain cancer | miR-34a-5p, miR-21-5p, miR-200a-3p, let-7b-5p |
| Head and neck cancer | miR-124-3p, miR-21-5p, miR-155-5p, miR-205-5p, miR-145-5p |
| Eye cancer | miR-18a-5p, miR-17-5p, miR-34a-5p |
| Thyroid cancer | miR-146-5p, miR-125a-5p, miR-21-5p, let-7 family (5p arm), miR-99b-5p, miR-145-5p |
| Lung cancer | let-7a-5p, miR-218-5p, miR-138-5p, miR-145-5p, miR-195-5p, miR-21-5p |
| Breast cancer | miR-21-5p, miR-200c-3p, miR-155-5p, miR-200b-3p |
| Kidney cancer | miR-200c-3p, miR-21-5p, miR-19a-3p |
| Bladder cancer | miR-145-5p, miR-125b-5p, miR-143-3p, miR-124-3p |
| Esophageal cancer | miR-21-5p, miR-145-5p, miR-125a-5p, miR-34a-5p |
| Stomach cancer | miR-21-5p, miR-19a-3p, miR-145-5p, miR-20a-5p |
| Colon cancer | miR-21-5p, miR-200c-3p, miR-200b-3p |
| Liver cancer | miR-21-5p, miR-221-3p, miR-106b-5p, miR-101-3p, miR-203, miR-34a-5p |
| Pancreatic cancer | miR-21-5p, miR-221-3p, miR-200c-3p, miR-34a-5p |
| Uterine cancer | miR-205-5p, miR-424-5p, miR-34a-5p, miR-101-3p, miR-200a-3p, miR-21-5p |
| Cervical cancer | miR-145-5p, miR-21-5p, miR-143-3p, miR-218-5p |
| Ovarian cancer | miR-21-5p, miR-145-5p, miR-205-5p, miR-125b-5p |
| Prostate cancer | miR-34a-5p, miR-205-5p, miR-143-3p, miR-195-5p |
| Blood cancer | miR-29b-3p, miR-21-5p, miR-15a-5p, miR-16-5p |
| Bone cancer | miR-34a-5p, miR-143-3p, miR-144-5p, miR-214-3p |
| Soft-tissue cancer | miR-520c-3p, miR-373-3p |
| Skin cancer | miR-211-5p, miR-21-5p, miR-205-5p, miR-203-3p |

A main feature of namiRa is the representation of miRNA functions as either oncogenic or tumor-suppressive. We also note that many miRNAs can have dual roles, not only in different contexts but also within the same context, depending on the specific genes targeted under particular cellular conditions [36]. Therefore, although the functional role of some miRNAs is well-established due to a high volume of publications, further investigation is required to clarify the interactions between other miRNAs and cancers with greater accuracy. Furthermore, we report validated target genes within the specific context in which a miRNA was investigated, aiming to uncover the genes mediating its functional mechanism. This contrasts with other databases, such as miRTarBase and TarBase, which catalog a large number of validated target genes without direct reference to the underlying functional mechanisms [15, 29].

We also provide a dynamic network diagram illustrating miRNA roles and their target genes, allowing researchers to quickly grasp the key insights into miRNA functional mechanisms. The combination of these features (*i.e.*, context-specific functional annotations and interactive visual networks) distinguishes namiRa from existing miRNA-cancer databases. For example, while miRCancer catalogs miRNA expression patterns, it lacks detailed functional relationships and cell line-specific resolution [13]. Similarly, miR2Disease provides miRNA-disease associations but offers limited depth for cancer-specific relationships and covers far fewer cancer types, encompassing only 349 miRNAs across 163 diseases [25]. Other resources, such as CMC, are primarily bioinformatic compendia listing cancer-related miRNAs without the rich experimental context [27], and dbDEMC similarly relies heavily on computational predictions [26]. OncomiRDB, while including experimentally verified miRNAs, has not been updated since 2014 and contains a limited set of only 328 entries [28]. Crucially, none of these databases provide a significant volume of experimentally validated regulatory mechanisms paired with a dynamic visual network to intuitively summarize miRNA interactions and functions from the literature. It is this integration of comprehensive, curated experimental data with advanced visualization tools that enables namiRa’s unique utility, allowing researchers to perform cross-cancer comparisons of individual miRNAs or analyze multiple miRNAs within a specific cancer, all while facilitating exploration of miRNA-target and miRNA-pathway associations through direct hyperlinks. Overall, namiRa addresses a critical gap in miRNA research by providing a centralized, high-quality resource that integrates dispersed data into a unified platform, facilitating the efficient exploration of miRNA dysregulation in cancer.

## Conclusion and future directions

namiRa is a comprehensive, manually curated database dedicated to cataloging miRNA-cancer associations. By systematically reviewing over 9,884 publications, we have compiled detailed information on 1,078 miRNAs across 33 specific cancer types. This includes data on miRNA expression patterns, functional roles, experimentally validated targets, and regulatory networks, all accessible through user-friendly graphical and text-based interfaces.

The database is freely accessible via a web interface and will be updated annually, with each new release incorporating additional miRNA-cancer relationships. While the current version of namiRa has been curated with reasonable accuracy, its comprehensiveness can be further improved by including new and previously overlooked publications on miRNAs and additional cancer types. Future development will also focus on integrating more online tools and data sources to enhance its utility.

Consequently, subsequent work will prioritize expanding the functionality of namiRa by increasing coverage of neglected cancers, refining the annotation of experimental validation data, and linking miRNA interactions to clinical outcomes. These enhancements will strengthen the applicability of namiRa in cancer research, precision oncology, and tumor biomarker discovery. As miRNA therapeutics advance, the curated insights provided by namiRa will serve as a valuable foundation for translational research, fostering breakthroughs in cancer diagnosis, prognosis, and targeted therapy. We anticipate that namiRa will aid the biomedical community in achieving a holistic understanding of miRNA function in cancer and accelerate the development of novel cancer therapeutics.

## Author contributions

S.Moradi: conceptualization, study and database design, funding acquisition, project administration, supervision, writing – review & editing, final approval of the manuscript. S.Mohebbi and S.A.: data acquisition and curation, validation, writing – original draft.

A.D.: database development, visualization.

Z.A., H.T., M.G., M.R., A.Y., N.R., F.M., A.A., S.Mosanan, F.A., L.A., R.F., N.S., H.S.: data acquisition, investigation.

## Data availability

The namiRa database will be continuously maintained and updated. The database is now publicly accessible at https://www.namira-db.com.

## Acknowledgments

The authors thank the Moradi lab members particularly Parisa Torabi for discussions and assistance with data acquisition. This work received no external funding.

